# Role of Toll-like receptor 2 (TLR2) in muscle fatigue: significance for muscle myopathies

**DOI:** 10.64898/2026.11.16.766957

**Authors:** Mei Bruton, Arthur Cheng, Kent Jardemark, Ghada Nouairia, Eric Rullman, Abram Katz, Niklas Ivarsson, Ingrid E. Lundberg, Håkan Westerblad, Helena Erlandsson Harris, Joseph Bruton

## Abstract

We report that Toll-like receptor 2 (TLR2) modulates skeletal muscle function. TLR2 ligands accelerated muscle fatigue development in isolated muscle fibers. Muscle fibers from TLR2 gene knock-out (TLR2-/-) mice were more fatigue resistant. This was not due to altered mitochondrial oxidative capacity or glycogen content. TLR2-/- muscle fibers had larger tetanic [Ca^2+^]_i_ transients. Muscles from TLR2-/- mice had increased levels of Ca^2+^-binding proteins compared to WT muscles. Lastly, TLR2 was expressed in muscle fibers of control subjects and patients with idiopathic inflammatory myopathies (IIM), whereas high mobility group box protein (HMGB) 1, an endogenous activator of TLR2, was only detected in the cytoplasm of muscles from IIM patients. In conclusion, TLR2 signaling regulates force production in skeletal muscle. TLR2 and its endogenous ligand, HMGB1 likely contribute to the muscle weakness in IIM and, possibly, other muscle disorders. Therapies targeted to the TLR2 signaling pathway may counteract pathological muscle weakness.

## Introduction

The Toll-like receptor 2 (TLR2) is a member of the Toll-like receptor family and is a transmembrane protein abundantly expressed in certain peripheral blood leukocytes (1, 2) and plays a role in the recognition of pathogen-associated molecular patterns (PAMPs), such as lipopeptides from bacteria (3). Activation of TLR2 by PAMPs initiates innate immune responses, triggers intracellular signaling pathways via myeloid differentiation primary response protein 88 (MyD88) and leads to the production of cytokines and chemokines. Thus, TLR2 stands in the frontline of a host’s defenses against pathogens and plays an important role in both innate and adaptive immunity (4–6).

Additional physiological roles for TLR2 can be envisioned. First, TLR2 is expressed not only in immune cells but also in a variety of other tissues including skeletal muscle (7, 8). Second, while TLR2 has a well-defined role in recognizing bacterial fragments, other ligands such as serum amyloid A, damage-associated molecular patterns (DAMPs) and high mobility group protein box 1 (HMGB1) activate TLR2 (3, 9–13). Third, genetic ablation or inhibition of TLR2 conferred a degree of protection against the metabolic dysfunction induced in skeletal muscle by high-fat feeding (14–16) and reduced infarct size in cardiac muscle (17, 18).

Currently, it is unknown whether TLR2 and its associated signaling pathways modulate skeletal muscle function. In the present study, we hypothesized that ablation of TLR2 or exposure to TLR2 ligands would impact skeletal muscle contractile and metabolic function.

## Results

### Both exogenous and endogenous ligands of TLR2 accelerate muscle fatigue in WT skeletal muscle fibers

The expression of TLR2 in skeletal muscles from WT mice was confirmed by immunohistochemical staining of muscle cross-sections. TLR2 was expressed mainly on the muscle sarcolemma (Figure 1 A, B). This finding was confirmed by immunofluorescence staining performed on enzymatically dissociated fibers from the flexor digitorum brevis (FDB) muscles of WT and TLR2-/- mice. TLR2 was expressed on the sarcolemma of WT (Figure 1C) but was completely absent in TLR2-/- muscle fibers (Figure 1D).

**Figure 1.**
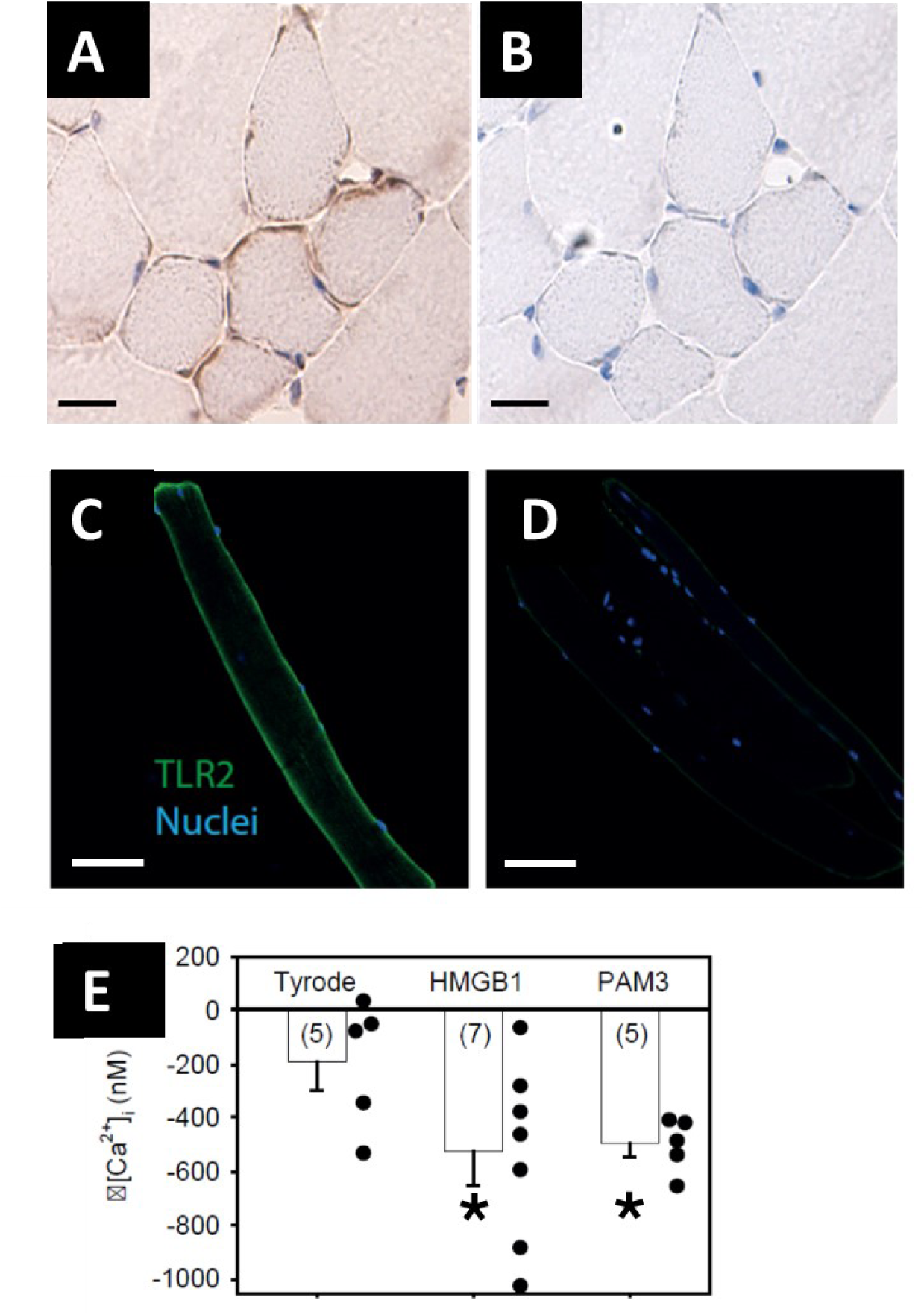
TLR2 is expressed in WT but not TLR2-/- muscle and its expression has a significant impact on muscle fatigue. **(A)** Immunohistochemical staining of muscle cross-sections shows that TLR2 is expressed in WT EDL muscle fibers, mainly on the membrane but also in the myoplasm. **(B)** Isotype staining as a negative control for TLR2. Images in **(A)** and **(B)** are representative of three independently performed experiments in four WT mice. Brown indicates positive staining and blue shows the nuclei. Scale bar is 25 µm. **(C, D)** Confocal immunofluorescence images confirm the predominant membrane expression of TLR2 protein in FDB muscle fibers from WT **(C)** but not from TLR2-/- **(D)** mice. The results are representative of fibers from three mice in each group. Green indicates TLR2 positive staining and blue indicates the nuclei. Scale bar 50µm**. (E)** Both endogenous (HMGB1) and exogenous (PAM3) ligands of TLR2 accelerated muscle fatigability in WT skeletal muscle fibers. Fatiguability was assessed as Δ[Ca^2+^]_i_ which is the change in [Ca^2+^]_i_ between the first and last tetanic stimulation. The graph shows clearly that both PAM3 and HMGB1 caused a significant decrease in tetanic [Ca^2+^]_i_ in FDB muscle fibers compared to fibers incubated in Tyrode only. The numbers in parentheses show the number of fibers; * indicates p value < 0.05. Filled circles show individual values.

Dissociated FDB muscle fibers were incubated with either HMGB1 (endogenous ligand) or Pam3CSK, a synthetic lipopeptide that mimics bacterial lipopeptides (exogenous ligand), for 12 hours. Control FDB fibers were exposed to the same conditions without any added ligands. After 12 hours, fibers were subjected to a fatigue protocol consisting of a series of 300 intermittent tetanic contractions. The index of fatigue was the decline in tetanic myoplasmic free calcium ([Ca^2+^]_i_) between the first and last of the 300 tetanic contractions (19). The addition of either HMGB1 or PAM3CSK caused a significantly greater decrease in tetanic [Ca^2+^]_i_ than under control conditions (Figure 1E). Collectively, these data indicate that TLR2 signaling exists in skeletal muscles and may be activated under conditions, such as infection and inflammation, where TLR2 ligands are generated.

### Skeletal muscle fibers from TLR2-/- mice show increased fatigue-resistance and greater tetanic [Ca^2+^]_i_

The above results show that activation of TLR2 by its ligands impair muscle function. Therefore, we compared function in muscle fibers from TLR2 gene knockout mice (TLR2-/-) with those from wild-type mice of the same genetic background (WT).

Single mechanically dissected FDB muscle fibers from C57BL/6 and TLR2-/- mice were fatigued with a protocol of repeated intermittent tetanic contractions (70 Hz, 350 ms duration at 2 s intervals) until tetanic force declined to 40%. Figure 2A shows that a significantly greater number of tetani were needed to reduce force to 40% in TLR2-/- fibers than in fibers from WT mice i.e. TLR2-/- fibers were more fatigue-resistant. Tetanic [Ca^2+^]_i_ was significantly greater in the first tetanus of the fatigue protocol in TLR2-/- than in WT muscle fibers but both decreased to the same level in the last tetanus of fatigue (Figure 2B). Tetanic force was similar in the TLR2-/- and WT fibers at the start and at the completion of the fatigue protocol (Figure 2E). Representative examples of tetanic [Ca^2+^]_i_ (Figure 2C and 2D) and force (Figure 2F and 2G) during the induction of fatigue emphasize that development of fatigue was delayed in TLR2-/- compared to WT fibers. Finally, we assessed whether fatigue resistance in TLR2-/- fibers was a consequence of the acute loss of functional TLR2. FDB fibers from WT mice were cultured overnight without or with 10 µM CU-CPT22, an inhibitor of TLR2 receptors. The index of fatigue was similar in the two situations and tetanic [Ca^2+^]_i_ at the end of 300 tetanic contractions was 0.80 ± 0.07 µM (n = 14) and 0.85 ± 0.09 µM (n=18) in the absence and presence of CU-CPT22 respectively. These findings indicate that the increased fatigue resistance in TLR2-/- fibers required long-term adaptations coupled to the loss of functional TLR2 receptors.

**Figure 2.**
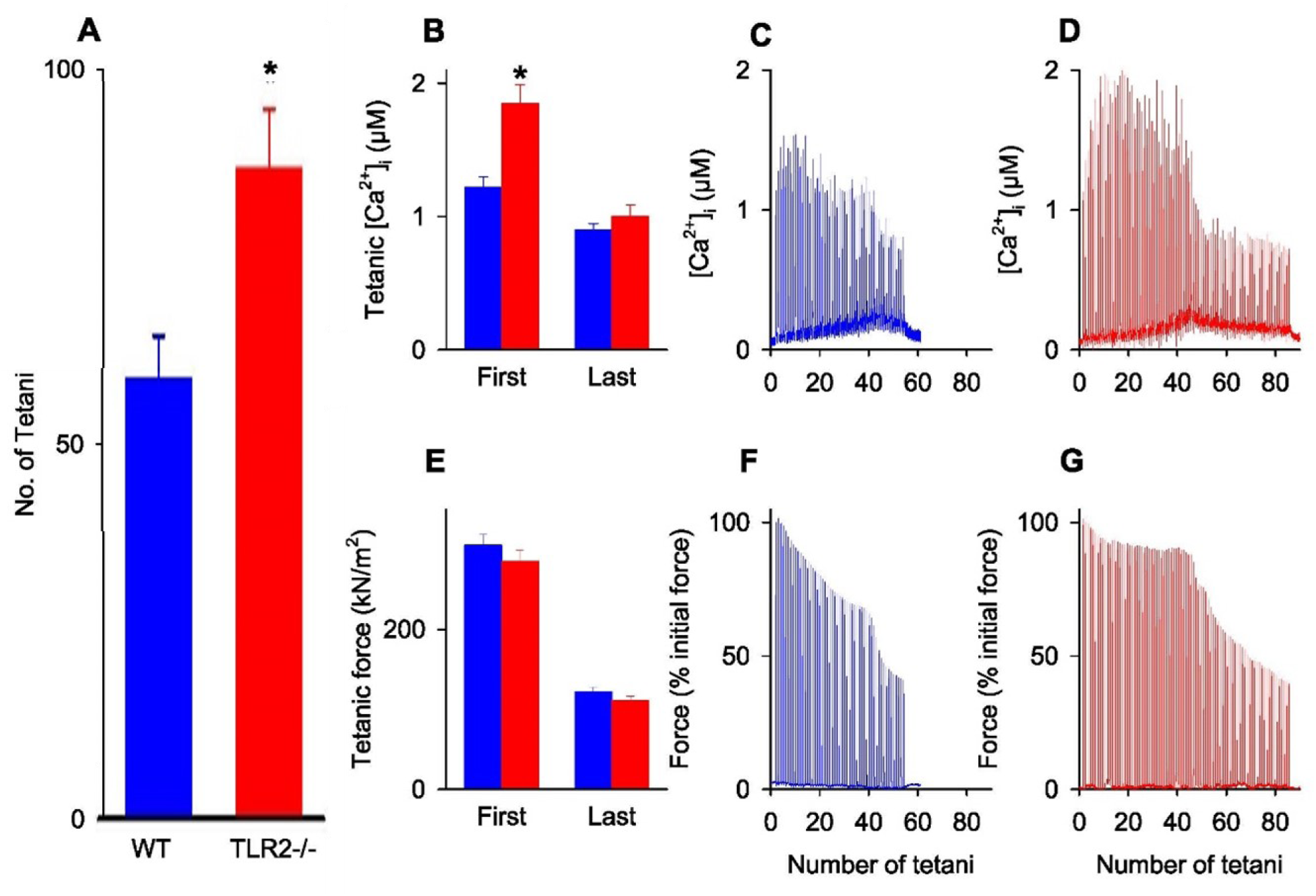
Skeletal muscle fibers from TLR2-/- mice were more resistant to fatigue compared to WT fibers. **(A)** The number of tetani needed to reduce tetanic force to 40% was significantly greater in TLR2-/- (red bars) than control (blue bars) fibers. **(B, C, D)** At the start of the fatigue protocol, the tetanic [Ca^2+^]_i_ in first tetanus was significantly greater in TLR2-/- than in WT fibers whereas it was similar in TLR2-/- and WT fibers in the last tetanus of the fatigue induction protocol. Typical tetanic [Ca^2+^]_i_ records during the fatigue protocol are shown in **C** (WT) and **D** (TLR2-/-). **(E, F, G)** Tetanic force generated by WT and TLR2-/- muscle fibers was similar at the start of the fatigue protocol and both decreased to the same extent by the end. Values in A, B, and E represent mean ± SEM, n = 20, * p value < 0.05. Typical records of tetanic force in a WT fiber and TLR2-/- fiber during induction of fatigue. are shown in **(F)** and **(G)**, respectively.

Next, we performed biochemical analyses to assess whether improved mitochondrial function could explain the increased fatigue resistance of TLR2-/- muscle fibers. We examined extensor digitorum longus (EDL) muscles which have the same fiber-type composition as the FDB muscles and were obtained from the same mice as the FDB muscles. However, we found no statistically significant differences in citrate synthase (CS, Figure 3A) and 3-hydroxyacyl-CoA dehydrogenase (HAD, Figure 3B) activity, or cytochrome c oxidase I (COX1, Figure 3D) protein expression between muscles from TLR2-/- and WT mice. Similarly, there was no significant difference between groups for glycogen, the major energy substrate during most forms of muscle contraction/exercise (Figure 3C). Finally, we explored whether muscle antioxidant capacity was altered in TLR2-/- mice by measuring superoxide dismutase (SOD) 1 and SOD2 protein expression. We found that in TLR2-/- mice, both SOD1 (Figure 3E) and SOD2 (Figure 3F) were elevated significantly compared with WT mice. Collectively, these data provide a molecular signature of muscles from TLR2-/- mice which do not explain the cause of the increased muscle fatigue resistance.

**Figure 3.**
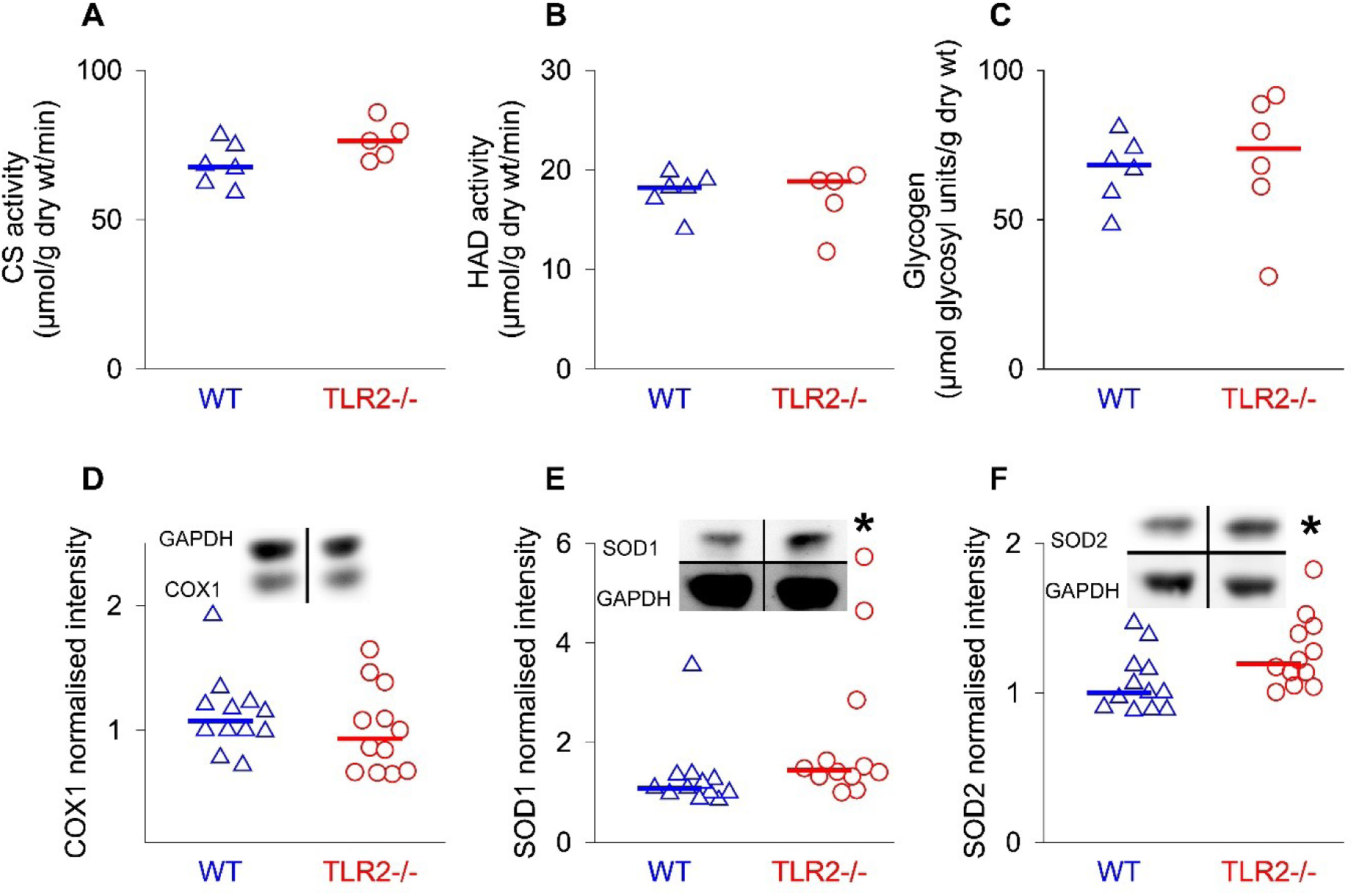
Mitochondrial energy-generating capacity and muscle glycogen were not changed but antioxidant capacity was increased in TLR2-/- compared to WT mice. Muscle mitochondrial energy-generating capacity was investigated by measuring (A) citrate synthase (CS) and (B) 3-hydroxyacyl-CoA dehydrogenase (HAD) activities with spectrophotometry and (D) cytochrome c oxidase I (COX1) with western blot. No significant differences were found in these proteins or glycogen content (C) in muscles from TLR2-/- compared to WT mice. Both cytosolic (SOD1, E) and mitochondrial (SOD2, F) superoxide dismutase protein expression were significantly elevated in muscles from TLR2-/- mice compared with WT mice, *p value < 0.05, Mann-Whitney rank sum test. All results are representative of three independently performed experiments in a total of at least five mice in each group. Each individual value is plotted as a separate symbol together with the median.

Tetanic force is determined by Ca^2+^ released from the sarcoplasmic reticulum (SR) of skeletal muscle fibers. We observed that while tetanic force at each stimulation frequency was similar in TLR2-/- and WT muscle fibers (Figure 4A), tetanic [Ca^2+^]_i_ was significantly higher at stimulation frequencies of 50 Hz and higher in TLR2-/- muscle fibers (Figure 4B). To examine why TLR2-/- mice had higher tetanic [Ca^2+^]_i_, we investigated two important Ca^2+^-binding proteins expressed in the SR: calsequestrin (Figure 4C) and sarcalumenin (Figure 4D) and found the levels of both to be significantly greater in TLR2-/- mice than in WT mice. We also investigated the proteins controlling Ca^2+^-release from the SR, the dihydropyridine receptor (DHPR) and ryanodine receptor (RyR). Neither DHPR (Figure 4E) nor RyR (Figure 4F) showed any significant difference in protein expression between TLR2-/- mice and WT mice. Thus, these data indicate that there is a higher content of Ca^2+^-buffering proteins in SR in skeletal muscle of TLR2-/- mice which ensures that bigger SR Ca^2+^ stores are available for release and suggest a mechanism underlying the bigger tetanic [Ca^2+^]_i_ transients observed during electrical stimulation.

**Figure 4.**
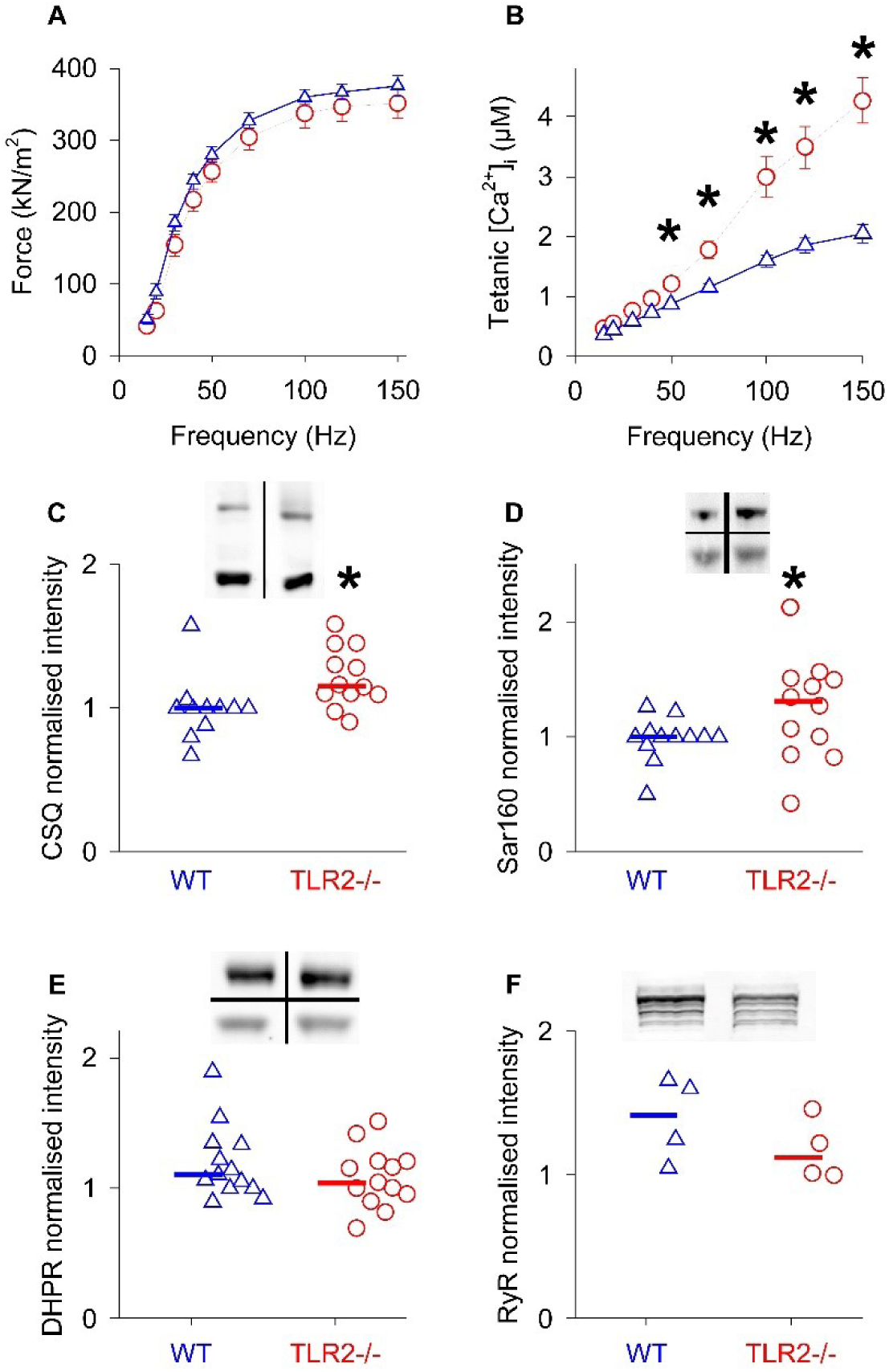
Force production is the same but tetanic [Ca^2+^]_i_ is greater in TLR2-/- than in WT muscle fibers. (A) At all stimulation frequencies, tetanic [Ca^2+^]_i_ is significantly greater (p < 0.001, 2-way ANOVA) in TLR2-/- (red circles) compared to WT muscle (blue triangles) while (B) tetanic force at all stimulation frequencies is similar in TLR2-/- and WT muscles. The sarcoplasmic reticulum calcium-binding proteins (C) calsequestrin (CSQ) and (D) sarcalumenin (Sar160) are significantly higher (*p < 0.05, Mann-Whitney rank sum test) in muscles from TLR2-/- mice than from WT mice. There was no difference in the protein expression of (E) dihydropyridine receptor (DHPR) or (F) ryanodine receptor (RyR) in muscles from TLR2-/- mice compared to those from WT mice. In C – F, representative western blots are shown on each panel and the results are representative of four independently performed experiments in a total of 12 mice in each group except for E and F where the results are representative of three independently performed experiments in a total of at least four mice in each group. In (A) and (B) values are shown as mean ± SEM (n = 20) while in (C – F) each individual value is shown as a separate symbol together with the median.

### Gene expression

Gene ontology (GO, 20, 21) and Kyoto Encyclopedia of Genes and Genomes (KEGG, 22) analyses were employed to investigate transcriptional alterations in skeletal muscle tissue from TLR2-/- mice. GO provides a structured framework to annotate gene function across species, encompassing biological processes, cellular components, and molecular functions. KEGG offers manually curated pathway maps integrating metabolic and signaling networks. In this study, a total of 8,486 probe sets were identified in the muscle samples. Functional enrichment analysis using GO and KEGG pathways was performed to characterize the molecular pathways associated with TLR2 deficiency (Figure 5A and 5B respectively). In particular, TLR2-associated signaling pathways were probed using publicly available pathway annotation databases to explain the downstream effects of TLR2 ablation.

**Figure 5.**
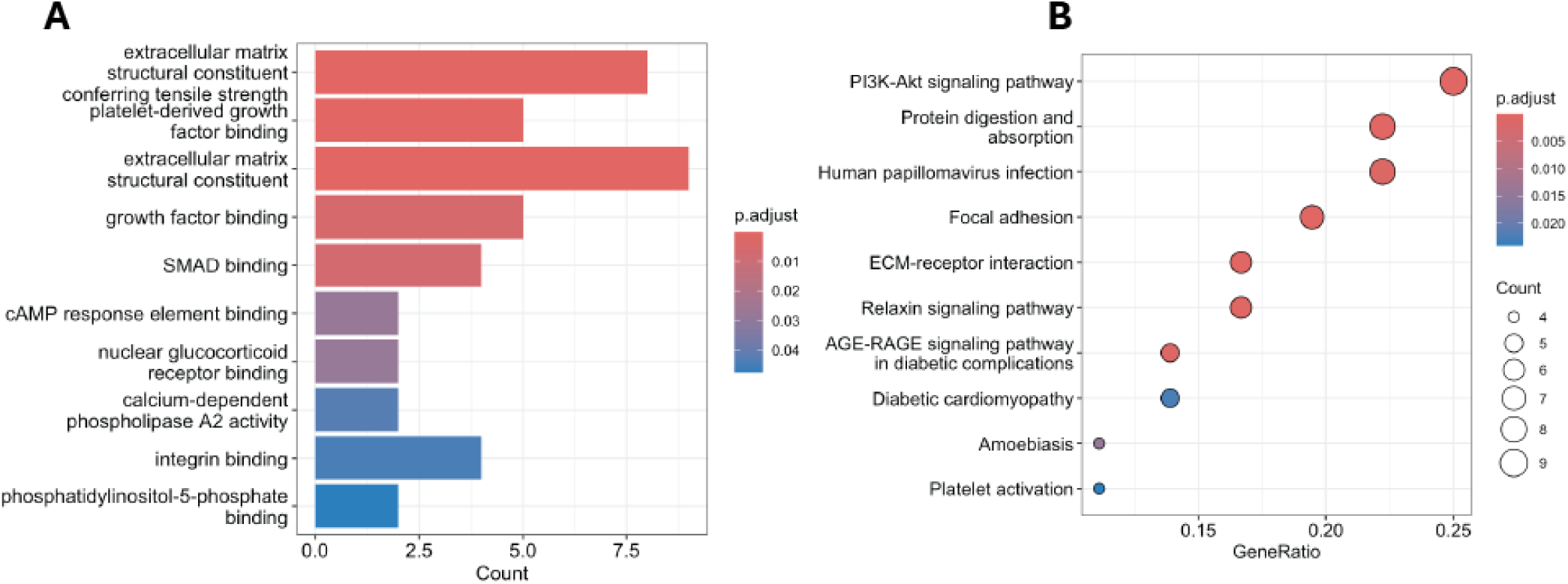
Functional enrichment analysis reveals extracellular matrix remodeling and growth factor signaling as key features in TLR2-/- mice. (A) Gene Ontology (GO) enrichment analysis of molecular functions among the significantly altered proteins highlights an overrepresentation of terms related to extracellular matrix (ECM) structural components, growth factor binding, and SMAD signaling. These functions are indicative of active tissue remodeling and TGF-β-related signaling, consistent with tumor microenvironment reprogramming. The bar plot shows the count of proteins annotated for each term, with color indicating adjusted p-values (Benjamini-Hochberg correction). (B) KEGG pathway enrichment analysis underscores the involvement of several oncogenic and microenvironment-related pathways, including PI3K-Akt signaling, ECM-receptor interaction, focal adhesion, and platelet activation. The x-axis represents the gene ratio (number of hits divided by the total input size), while dot size reflects the number of proteins per pathway and dot color encodes statistical significance. These pathways suggest activation of survival, adhesion, and inflammation-related mechanisms that may underlie tumor progression or stromal interactions.

Among the top enriched pathways were Toll-like receptor signaling, NF-κB signaling, and cytokine-cytokine receptor interaction, highlighting the immunomodulatory role of TLR2. In line with its known function, TLR2 deficiency was associated with downregulation of key inflammatory mediators and signal transducers, including *Myd88*, *Il6*, and *Tnf*. Additional enrichment was observed in pathways related to muscle regeneration, apoptotic signaling, and oxidative phosphorylation, suggesting broader systemic effects of TLR2 deletion beyond innate immunity.

Collectively, these data indicate that loss of TLR2 signaling significantly remodels transcriptional and pathway landscapes in muscle tissue, with implications for inflammation, regeneration, and systemic energy homeostasis.

### TLR2 and HMGB1 are detected in skeletal muscles of patients with idiopathic inflammatory myopathies

Myositis or idiopathic inflammatory myopathies are rare autoimmune conditions that target skeletal muscles as well as other body systems and cause severe muscle fatigue and muscle weakness in patients. Inflammatory mononuclear cell infiltrates can be found in the muscles of patients and are regarded as a characteristic histopathological feature. Furthermore, the myocytotoxic effects of T cells that result in muscle necrosis are regarded as a likely mechanism leading to muscle weakness (23–25). However, many patients reporting muscle weakness and fatigue show no detectable infiltrates in their muscle biopsies (26). Thus, additional mechanisms may underlie the muscle dysfunction in these patients. Based on the increased fatigue resistance of muscle fibers from TLR2-/- mice, we hypothesized that aberrant TLR2 signaling could be responsible for the muscle fatigue and weakness in patients with myositis. We investigated the expression of TLR2 in muscle biopsies of six healthy controls and six subjects with myositis. TLR2 was expressed in muscle tissues of patients with myositis, both in the muscle fibers and endothelial cells/capillaries and even in muscles where inflammatory cell infiltrates were undetectable (Figure 6A and inset). HMGB1 was found in the cytoplasm of endothelial cells and muscle fibers of these patients. In the healthy controls, we detected TLR2 expression in muscle fibers and endothelial cells/capillaries in five of the six subjects studied (Figure 6C and inset). It is noteworthy that cytosolic location of HMGB1 was never seen in endothelial cells or in muscle fibers obtained from the healthy subjects. Isotype negative samples are shown for comparison in Figure 6B and 6D. Collectively, these data suggest that it is the combination of TLR2 together with extra-nuclear located HMGB1 that contributes to the pathology of myositis.

**Figure 6.**
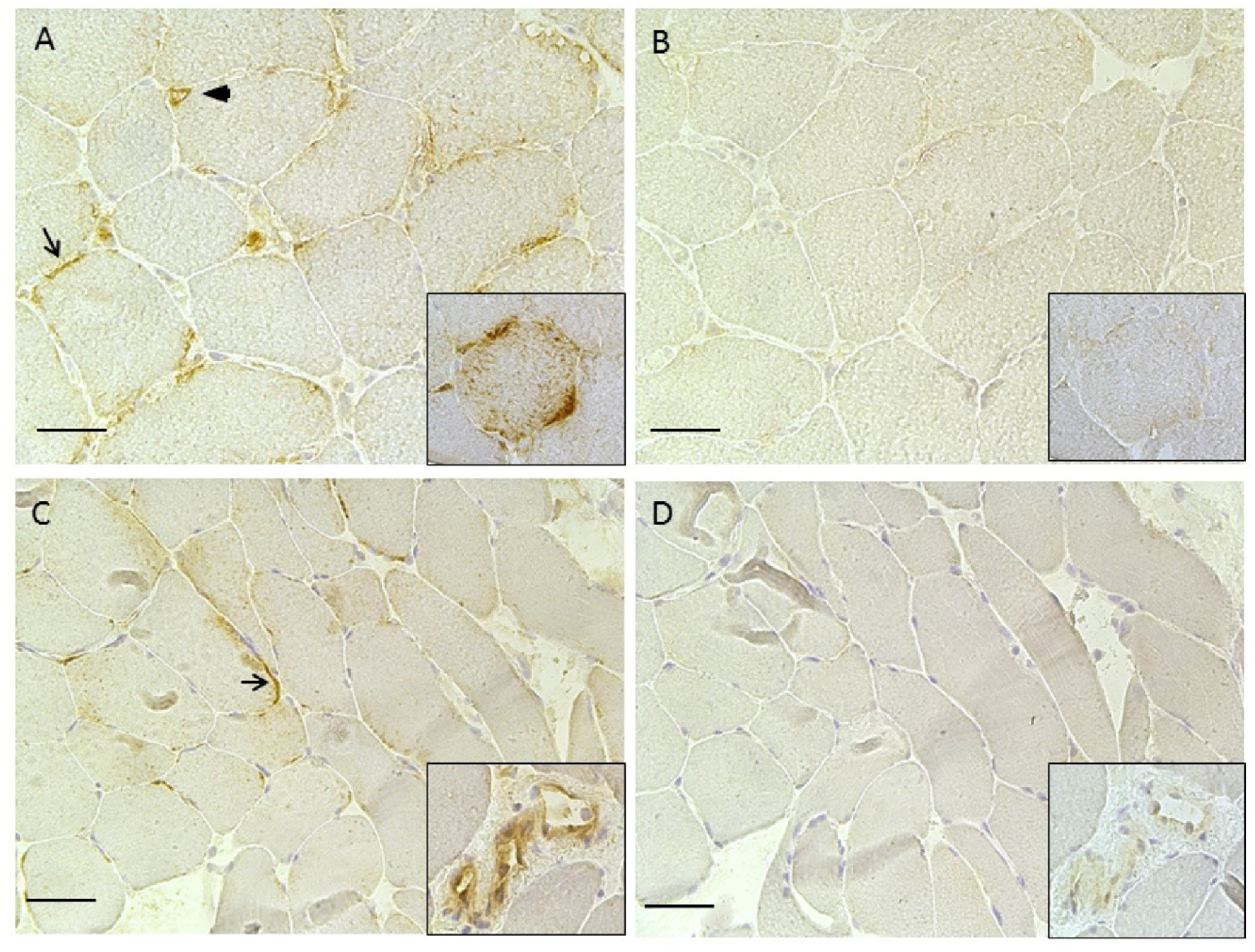
TLR2 is expressed in the muscles of patients with myositis and healthy individuals. (A) Representative immunohistochemical staining of TLR2 expression in the muscle of a patient with myositis. Arrow shows TLR2 expression on the membrane of a muscle fiber. Arrowhead shows TLR2 expression in a capillary. Inset shows a muscle fiber with TLR2 expression both on membrane and in the cytosol. (B) consecutive section from the same patient but stained with isotype as negative control for TLR2. (C) representative picture shows TLR2 expression in the muscle of a healthy individual. Arrow shows TLR2 expression on the membrane of a muscle fiber. Inset showed TLR2 expression on a large vessel. (D) a consecutive section from the same healthy individual but stained with isotype as negative control for TLR2. Brown colour indicates positive staining. Blue colour shows the nuclei. Magnification 20x. Scale bar 25µm.

## Discussion

The major novel findings in this study are: (1) TLR2 is expressed in adult skeletal muscles of C57BL/6 mice and muscle contractile function is depressed via a TLR2-activated pathway through infection (Pam3CSK) or inflammation (HMGB1) and (2) muscle fatigue resistance is improved when TLR2 is absent from the muscles of mice.

In common with other TLRs family members, TLR2 expression in immune cells undertakes multiple roles in the immune response to infection. However, increasing numbers of studies show that TLR2 is present in non-immune tissues including skin, kidney, rectum and ovary (27, 28). The expression of TLR2 in skeletal muscle and its up-regulation in rodents in response to high-fat diets and insulin resistance as well as in mouse models of dystrophy was reported earlier (8, 29, 30) In the current study, we found that *in vitro* treatment of adult mouse skeletal muscle fibers with TLR2 agonists, such as Pam3CSK and HMGB1, accelerated the development of muscle fatigue. Pam3CSK is a synthetic triacylated lipopeptide that mimics the release of lipopeptide in bacterial infections, while HMGB1 functions as an alarmin that is released by damaged cells during sterile processes such as autoimmune disease (31). Thus, TLR2 activation in skeletal muscle might contribute to the marked sensation of muscle fatigue experienced by individuals with an infection or inflammation such as influenza. Our data suggest that this fatigue is mediated by activation of the TLR2 receptors that results in decreased levels of myoplasmic Ca^2+^. However, the specific mechanisms or pathways through which ligands act to accelerate Ca^2+^ depletion and muscle fatigue are unclear.

In the absence of TLR2 expression, we found that skeletal muscle fibers were more fatigue resistant. Surprisingly, this was not accompanied by any increase in mitochondrial enzyme activities or increased mitochondrial biogenesis in TLR2-/- muscle compared to WT muscles. Basal muscle glycogen content, depletion of which is implicated in the development of muscle fatigue, was not different between TLR2-/- and WT muscle. A more robust antioxidant capability was present in the muscles of the TLR2-/- mice as shown by the significantly increased levels of cytoplasmic SOD1 and mitochondrial SOD2 protein expression compared to the wild type mice. These findings are consistent with earlier studies showing that muscle weakness and accelerated fatigue developments are associated with SOD deficiency and/or increased superoxide (32–35) During the repeated intermittent tetanic contractions used to induce muscle fatigue, tetanic [Ca^2+^] and force generation declined more slowly in muscle fibers from TLR2-/- than WT mice. This is likely a consequence of the greater amounts of the SR Ca^2+^ storage proteins, calsequestrin and sarcalumenin (Figure 5), which results in more Ca^2+^ available for release from the SR.

Gene expression profiling revealed that a relatively limited number of genes were significantly altered in the skeletal muscle of TLR2⁻/⁻ mice. Notably, none of the differentially expressed genes encoded proteins were directly associated with energy metabolism or mitochondrial function, which are commonly affected in muscle pathologies (36, 37). Instead, the majority of altered transcripts corresponded to extracellular matrix (ECM) components, protein digestion and absorption, and elements of the PI3K-Akt signaling pathway.

The transcriptional profile of the TLR2-/- mice is in stark contrast to findings in muscular dystrophies and inflammatory myopathies, where genes encoding ECM proteins—especially collagens and matrix-remodelling enzymes—are typically upregulated, reflecting fibrotic and regenerative responses to muscle injury (38,39). In the TLR2-deficient muscle, however, these ECM-related genes were downregulated, suggesting a fundamentally different remodelling response in the absence of TLR2-mediated signaling. Interestingly, the PI3K-Akt pathway, which is also altered in TLR2⁻/⁻ mice, has been implicated in muscle hypertrophy (40). This implies that TLR2 may indirectly modulate structural adaptation and immune interactions through ECM remodelling and membrane protein expression, rather than directly altering energy metabolism.

The observed downregulation of ECM-related genes in TLR2⁻/⁻ muscle could reflect a blunted immune or fibrotic response, potentially offering a protective effect against excessive remodelling or inflammation. Given the importance of ECM composition and cell surface receptors in capillary distribution, ligand accessibility, and receptor activation, it is plausible that TLR2 plays a regulatory role at the interface of innate immunity and tissue architecture. Further studies are warranted to investigate how TLR2-dependent pathways intersect with structural remodelling and immune regulation in skeletal muscle.

Based on the present data, we conclude that the TLR2 signaling pathway plays an important role in skeletal muscle function. Therefore, modifying the TLR2 pathway could be a valuable treatment for patients with innate muscle dysfunction.

## Methods

### Mice and ethical approval

All experiment procedures complied with the Swedish Animal Welfare Act, the Swedish Welfare ordinance, and applicable regulations and recommendations from Swedish authorities. The study was approved by the Stockholm North local ethics committee on Animal Experiments. WT C57BL/6 and TLR2-/- mice were used in this study. TLR2-/- mice were on a C57BL/6 background and were kindly provided by Dr S. Akira (Osaka University, Osaka, Japan). WT C57BL/6 mice were purchased from Charles River Laboratories (Wilmington, MA, USA). All the mice used in this study were males and more than 7 weeks old. Mean body mass was not significantly different between the TLR2 -/- mice and the WT mice being 30.3 ± 0.6 g (n = 30) and 31.8 ± 0.6 g (n = 16) respectively. This is in concordance with previous reports of bodyweight in TLR2 -/- and WT mice (Zbinden-Foncea et al 2012; Kim el al 2015). .Animals were kept and bred at Karolinska Institutet animal facility in a specific pathogen-free environment with standard room temperature and a 12-12 hour light/dark cycle. Food and water were provided *ad libitum*. Mice were killed by rapid neck disarticulation and the FDB and EDL muscles were removed.

### Enzymatic single fiber dissociation, TLR2 ligands stimulation and [Ca^2+^]_i_ measurements

Enzymatically dissociated FDB muscle fibers were isolated as described previously (41). Briefly, the whole FDB muscles were removed from both hind paws of WT and TLR2-/- mice. The tendons, connective tissue and blood vessels around FDB muscles were removed under a microscope using a pair of forceps and micro-iris scissors. Clean FDB muscles were incubated for 2-3 hours at 37 °C in 0.3% collagenase type I (Sigma) in DMEM (Invitrogen) supplemented with 10% fetal bovine serum (Invitrogen). Muscles were transferred to fresh DMEM and gently triturated to dissociate individual muscle fibers. 300 µl of the resultant muscle fiber suspension was then transferred to laminin (Sigma) coated glass-bottom Petri dishes (Mattek) and allowed to attach for up to 15 minutes before 2.7 ml DMEM supplemented with antibiotic, antimycotic solution (1 μL/ml, Sigma) was added.

To investigate the effects of TLR2 ligands on tetanic [Ca^2+^]_i_, either Pam_3_CSK_4_ (PAM3, InvivoGen, San Diego, CA, USA) or HMGB1 was added to the solution bathing the FDB single fibers attached to the glass-bottom dishes with the final concentration being 10 µg/ml. CU-CPT22 (Tocris, Bristol, United Kingdom), a specific TLR1/2 antagonist (42) was added to other dishes with attached FDB single fibers (final concentration was 10 µM). Fibers were incubated at 37 ℃ for 12 h before measurement of tetanic [Ca^2+^]_i_. Control fibers were incubated under the same condition in the absence of ligands. The HMGB1 used in this study was endotoxin-free and disulfide-bonded (12, 43) and was provided by Dr. K. J. Tracey (Feinstein Institute for Medical Research, Manhasset, NY).

+

After 12 hours incubation of the WT FDB fibers with Pam_3_CSK_4_, HMGB1, or CU-CPT22 [Ca^2+^]_i_ was measured during a fatiguing protocol consisting of 300 intermittent tetanic contractions (70 Hz, 350 ms duration, 2 s intervals). [Ca^2+^]_i_ was measured in dissociated FDB fibers using the fluorescent Ca^2+^ indicator indo-1 (Invitrogen) as described previously (17). Briefly, muscle fibers attached to laminin-coated glass-bottom dishes were incubated with indo-1 AM (4 µM) for 20 min, followed by a 20 min wash. Change in indo-1 fluorescence was measured with a PTI system (Horiba) consisting of a Xenon lamp, a monochromator and two photomultiplier tubes. Excitation light was 360 ± 5 nm and the emitted fluorescence was collected after passage through a dichroic mirror at 405 ± 5 nm and 495 ± 5 nm. The ratio of the light emitted at 405 nm and 495 nm was translated to [Ca^2+^]_i_ as described previously (44).

### Immunofluorescence staining of TLR2 on FDB fibers from WT and TLR2-/- mice

Dissociated FDB fibers from WT and TLR2-/- mice without exposure to any ligands were incubated for 12 hours at 37°C. The medium was then removed and the fibers washed with cold PBS twice and fixed with 50% cold acetone for 30 seconds and with 100% cold acetone for 3 minutes as described previously (19). Muscle fibers were immediately stained for immunofluorescence according to a standard protocol (45). The primary antibody used for TLR2 identification was a purified goat polyclonal anti-TLR2 antibody (2.5 µg/ml, clone N-17, catalog sc-8689, Santa Cruz Biotechnology Inc). The secondary antibody was a biotinylated donkey anti-sheep/goat IgG (AB360, The Binding Site). Isotype control was a goat gamma globulin (34764, Jackson ImmunoResearch). Fluorophore was Avidin Oregon Green (Invitrogen, Eugene, OR, USA). Images of longitudinal sections of stained intact single muscle fibers were obtained with confocal microscopy using a Bio-Rad MRC 1024 confocal unit with a dual Calypso laser (Cobolt, Solna, Sweden) attached to a Nikon Diaphot 200 inverted microscope and a Nikon Plan Apo 40x oil-immersion objective, N.A. 1.3 (Nikon, Tokyo, Japan). Excitation was at 491 nm and the emitted light was collected through a 522 nm narrow band pass filter.

### Mechanical single fiber dissection, force and [Ca^2+^]_i_ measurements

Mechanically dissected single fibers were used in this study to measure muscle force and [Ca^2+^]_i_. Single fibers were isolated under a dark-field illumination microscope using a pair of forceps and micro-iris scissors. The FDB muscles were split into three bundles after which each bundle was trimmed down to two or three fibers. One fiber was selected and the rest were killed by cutting through the sarcolemma, leaving only the selected fiber viable. The fiber’s ability to contract was tested by focal electrical stimulation. After isolation of a single fiber, the tendons were gripped by platinum-foil micro-clips and the preparation was transferred to the perfusion channel of an experimental chamber, placed on the stage of an inverted microscope. The bottom of the chamber consisted of a thin glass cover slip and fibers were mounted as close as possible to this. The fiber was flanked by platinum electrodes which were used for stimulation.

Force from mechanically dissected fibers was measured using an Akers 801 force transducer vertically mounted in the experimental chamber. The force transducer was provided with a glass tube extension with a fine platinum hook at the end. On the other end of the chamber, there was an adjustable hook. The tendon clips were attached to the two hooks, suspending the fiber horizontally. The fiber was stretched to the length at which maximal tetanic force was produced and the diameter of the fiber at this length was measured in order to calculate the developed force per cross-sectional area. The cross-sectional areas of TLR2-/- and WT muscle fibers were not significantly different (862 ± 17 µm^2^ (n = 17) and 994 ± 25 µm^2^ (n = 20), respectively).

The [Ca^2+^]_i_ measurements in these dissected fibers were obtained in the same manner as for the enzymatically dissociated fibers described above with one minor difference. The dissected fibers were loaded with indo-1 by pressure-injecting the potassium salt of indo-1 into the fibers.

### Western blot, spectrophotometry and fluorometry

The EDL muscles were homogenized with a motor-driven ground glass homogenizer in ice cold homogenization buffer (20 µl per mg wet weight; pH 7.4) consisting of (mM): 20 Hepes, 150 NaCl, 5 EDTA, 25 NaF, 1 Na_3_VO_4_, 20% glycerol (v/v), 0.5% Triton X-100 (v/v), 1 tablet protease inhibitor cocktail (Roche, Basel, Switzerland) per 50 ml. The homogenate was centrifuged at 700g for 5 min at 4°C. The supernatant was transferred to a new tube and the protein concentration was determined using the Bradford assay (Bio-Rad, Hemel Hemstead, UK). Lysates were diluted 1:1in Laemmli buffer (Bio-Rad, Hercules, CA, USA), left overnight at 4 °C and then heated for 10 min at 70 °C. Subsequently, 10 µg protein was loaded and run on a NuPAGE Novex 4-12% precast Bis-Tris gels (Invitrogen). Proteins were then transferred to polyvinylidine fluoride membranes (Immobilon FL; Millipore, Billerica, MA, USA). Membranes were blocked for 1 hour at room temperature in blocking buffer (Li-COR Biosciences, Lincoln, NE, USA), followed by incubation overnight at 4 °C with anti-copper/zinc superoxide dismutase (anti-SOD1) (1:1000; Abcam, Cambridge, UK), anti-manganese SOD (anti-SOD2) (1:1000; Upstate Biotechnology, Lake Placid, NY), anti-OxPhos Complex IV subunit I (anti-COX) (1:1000, Invitrogen, USA), anti-sarcalumenin (1:1000, XIIC4, Abcam), anti-calsequestrin (1:1000, VIIID12, Abcam), anti-RyR (1:1000, Abcam), anti-dihydropyridine receptor (1:500, Abcam), anti-actin (1:1000; sigma) and anti-GAPDH (glyceraldehyde-3-phosphate dehydrogenase, 1:1000, Catalog No. 39-8600, Invitrogen, CA, USA). Membranes were washed in Tris-buffered saline-Tween 20 (TBS-T) and incubated for 1 hour at room temperature with secondary antibody (IRDye 680-conjugated goat anti-mouse IgG, IRDye 800CW or anti-Rabbit IgG, IRDye 680CW,1:15000, Li-COR Bioscience). Membranes were then washed in TBS-T and immunoreactive bands were visualized by an infrared fluorescence microscopy (IR-Odyssey Scanner, LI-COR Biosciences). Images were quantified with ImageJ. Density values for above markers were normalized with values for GAPDH for each muscle sample.

To measure activities of the enzymes citrate synthase and HAD, EDL muscles were freeze dried, dissected free of non-muscle constituents, powdered, thoroughly mixed and homogenized (25 µl/mg dry wt) with ground glass homogenizers in ice-cold buffer consisting of (in mM) 50 Tris, 5 citrate, 0.6 MnCl_2_, and 1 cysteine, with 0.05% (vol/vol) triton X-100, pH 7.4 (46). The homogenate was centrifuged at 1.400g at 4°C for 1 min. Aliquots of supernatant were frozen for subsequent analyses of citrate synthase and HAD with standard spectrophotometric techniques (47, 48).

For analysis of glycogen, aliquots of powdered EDL muscles were digested with hot 1 N KOH and hydrolyzed enzymatically to free glucose. Glucose was then analyzed enzymatically with a fluorometric technique (49).

### Gene analysis

Total RNA was extracted from EDL muscles from TLR2-/- and WT mice using TRIzol reagent (Invitrogen, Life Technologies), following the manufacturer’s protocol. RNA quality and concentration were assessed with a NanoDrop 2000 spectrophotometer (ThermoFisher Scientific), and samples were dissolved in 20 µl of RNase-free water without additives. Transcriptomic analysis was performed using the Affymetrix Mouse Gene 2.1 ST microarray plate (ThermoFisher Scientific). Sample preparation, cDNA synthesis, hybridization, and scanning were conducted at the Bioinformatics and Expression Analysis (BEA) Core Facility, Karolinska Institutet, Huddinge Hospital, Sweden. cDNA was hybridized to the arrays and scanned using the GeneChip Scanner 3000 7G.

Raw intensity data (CEL files) were background-corrected and normalized using the Robust Multi-array Average (RMA) method. Quality control assessments were performed using principal component analysis (PCA) and normalized unscaled standard error (NUSE) plots, confirming consistent array performance and identifying no outliers. Differential expression analysis was conducted using the limma package v3.28.14 in R (50), applying empirical Bayes moderated *t*-statistics. Probe sets were considered differentially expressed at a Benjamini-Hochberg (51) adjusted p-value < 0.05.

To characterize the biological significance of these transcriptional changes, we employed functional enrichment analysis using both the GO and the KEGG databases. GO analysis categorizes differentially expressed genes into relevant biological processes, molecular functions, and cellular components, while KEGG pathway analysis maps genes to curated metabolic and signaling pathways. Enrichment analyses were performed using the DAVID platform and confirmed using the R clusterProfiler. A Benjamini-Hochberg correction was applied to control the false discovery rate (FDR), and pathways with an adjusted p-value (FDR) < 0.05 were considered statistically significant.

The complete raw and processed microarray data have been deposited in the Gene Expression Omnibus (GEO) under accession number GSE103337.

### Myositis patients, muscle biopsies and ethical approval

This study was approved by the Regional Ethical Review Board in Stockholm and all patients and control individuals gave their informed written consent to participate in the study. Six patients diagnosed with polymyositis or dermatomyositis (52, 53) with established disease and persisting muscle weakness despite conventional immunosuppressive treatment (5 women and 1 man; median age = 69.5 years, range 55-75; median disease duration=47 months, range 18 months-102 months) were included in this study. Their detailed clinical data have been described previously (54). They were selected for having few inflammatory cell infiltrates in their muscle biopsies. Muscle biopsy samples from six healthy, age-matched controls (4 women and 2 men, median age 68.5 years, range 42-77) without clinical or histopathological signs of any muscle disease were included as controls.

Muscle biopsies were obtained either from vastus lateralis or tibialis anterior by a “semi-open” muscle biopsy technique under local anesthesia (44). The biopsy samples were frozen in isopentane precooled by immersion in liquid nitrogen and then stored at -80 ℃ until cryostat sectioning was performed. At least two pieces of muscle tissues were taken from each subject at the same time. The first and last sections from each series of consecutive sections of the same patient were stained with Mayer’s hematoxylin and eosin to estimate inflammatory infiltrates. Neither section showed any obvious signs of cellular infiltrates.

### Immunohistochemistry staining in muscle biopsies of patients and mice

The muscle biopsy specimens were cut into 7 μm thick sections using a cryostat and embedded on chrome gelatin-coated glass slides. They were air dried completely and then put into cold 2% formaldehyde in PBS for fixation for 20 min in the dark. After fixation the slides were washed twice in cold phosphate buffered saline (PBS) for 5 minutes each time. The slides were then exposed to 0.1% saponin in PBS for 10 min followed by blocking with 1% H_2_O_2_ in PBS for 1 hour in darkness, then blocking with avdin/biotin for 30 min each, and finally blocked with 5% normal horse serum for HMGB1 or with normal goat serum for TLR2. For identification of HMGB1expression, sections were stained with a noncommercial monoclonal mouse anti-HMGB1 antibody (2 µg/ml, clone 2G7, produced-in house at the former Critical Therapeutics, Boston, MA, USA; now Cornerstone Therapeutics, Cary, NC, USA). For identification of TLR2, sections were stained with a rabbit anti-TLR2 antibody (1 µg/ml, ab24192, Abcam). As negative controls, we used irrelevant mouse IgG2b (2µg/ml, X0944, Dako) for HMGB1and rabbit immunoglobulin fraction (1 µg/ml, X0903, Dako) for TLR2. The sections were incubated with the above-mentioned antibodies or isotype negative controls diluted in 3% normal human serum overnight at room temperature in a humid chamber. They were washed and blocked with 5% serum again as above before applying secondary antibodies, biotinylated horse anti-mouse IgG (H+L) (1/320, BA-2001, Vector) for HMGB1 and biotinylated goat anti-rabbit IgG (H+L) (1/1600, BA-1000, Vector) for TLR2 for 30 min. After washing, sections were incubated with avidin-biotin-peroxidase complex (ABC elite, PK-6100, Vector) for 45 min. Reactions were developed using a peroxidase substrate kit (SK-4100, Vector) containing 3, 3’-diaminobenzidine, with Maye’s hematoxylin counterstaining. Slides were mounted using glycerol. Stained tissue sections were examined using a Polyvar II microscope (Reichert-Jung, Vienna, Austria) and a digital Leica 300F camera (Leica, Cambridge, UK).

### Statistical analysis

Statistical analyses were performed using Statistica V 10.0 from StatSoft (Tulsa, OK, USA) or Sigmaplot (Systat Software Inc., San Jose, CA, USA). Data are presented as mean S.E.M. Due to the small sample number of patients and the non-normality of the data set, these data were tested for significance with non-parametric tests. P values < 0.05 were considered to be statistically significant.

## Acknowlegments

This work was funded by the Swedish Research Council, King Gustaf V 80-year Foundation, Funds at Karolinska Institute, and Lars Hierta Minne Stiftelsen.

